# Proteomic Profiling of Mammalian COPII Vesicles

**DOI:** 10.1101/253294

**Authors:** Frank Adolf, Manuel Rhiel, Bernd Hessling, Andrea Hellwig, Felix T. Wieland

## Abstract

Intracellular transport and homeostasis of the endomembrane system in eukaryotic cells depend on formation and fusion of vesicular carriers. COPII vesicles export newly synthesized secretory proteins from the endoplasmic reticulum (ER). They are formed by sequential recruitment of the small GTP binding protein Sar1, the inner coat complex Sec23/24, and the outer coat complex Sec13/31. In order to investigate the roles of mammalian Sec24 isoforms in cargo sorting, we have combined *in vitro* COPII vesicle reconstitutions with SILAC-based mass spectrometric analysis. This approach enabled us to identify the core proteome of mammalian COPII vesicles. Comparison of the proteomes generated from vesicles with different Sec24 isoforms confirms several established isoform-dependent cargo proteins, and identifies ERGIC1 and CNIH1 as novel Sec24C‐ and Sec24A-specific cargo proteins, respectively. Proteomic analysis of vesicles reconstituted with a Sec24C mutant, bearing a compromised binding site for the ER-to-Golgi QSNARE Syntaxin5, revealed that the SM/Munc18 protein SCFD1 binds to Syntaxin5 prior to its sorting into COPII vesicles. Furthermore, analysis of Sec24D mutants implicated in the development of a syndromic form of osteogenesis imperfecta showed sorting defects for the three ER-to-Golgi QSNAREs Syntaxin5, GS27, and Bet1.

## Introduction

A hallmark of eukaryotic cells is the presence of an elaborate endomembrane system, which divides their interior into spatially and functionally separated compartments. Vesicular transport pathways connect these organelles with each other to maintain their characteristic compositions and ensure their specialized functions. Among the different vesicular coat systems the best characterised are Clathrin-coated vesicle (CCVs), COPI and COPII vesicles. CCVs operate in the late secretory/endocytic pathway, COPI vesicles are involved in retrograde transport from the ER-Golgi intermediate compartment (ERGIC) and cis-Golgi cisternae towards the endoplasmic reticulum (ER), as well as in intra-Golgi transport. COPII-coated vesicles promote anterograde ER-to-Golgi transport, the first step of newly synthesized secretory proteins and proteins destined for organelles other than the ER (Adolf and Wieland, 2014; Barlowe and Helenius, 2016; Bethune and Wieland, 2018; Miller and Schekman, 2013; Venditti et al., 2014).

The COPII coat, comprising the five cytosolic proteins Secretion-associated Ras-related protein 1 (Sar1), the Sec23/24 complex, and the Sec13/31 complex (Barlowe et al., 1994) is assembled on specialized subdomains of the ER, termed ER exit sites (ERES) or transitional ER (tER) (Bannykh et al., 1996; Orci et al., 1991; Rossanese et al., 1999). The biogenesis of COPII vesicle is initiated by recruitment and activation of Sar1 on ER membranes by the ER-resident guanine nucleotide exchange factor (GEF) Sec12 (Barlowe and Schekman, 1993; d’Enfert et al., 1991; Nakano et al., 1988). Nucleotide exchange within the small GTP binding protein triggers a conformational change resulting in the exposure of an amphipathic N-terminal helix by which Sar1 is anchored to the ER-membrane (Bi et al., 2002; Huang et al., 2001). Subsequently, the COPII coat is assembled by successive recruitment of the inner and outer coat subcomplexes (Matsuoka et al., 1998). Activated membrane-bound Sar1, by interaction with the Sec23 subunit, first recruits the heterodimer Sec23/24, to form the inner COPII coat layer (Bi et al., 2002). Seemingly paradoxical, the Sec23 subunit at the same time acts as GTPase activating protein (GAP) of Sar1 (Yoshihisa et al., 1993) and is further stimulated by binding of the outer coat subunit Sec31 (Antonny et al., 2001). The Sec24 subunit as the major cargo-binding subunit is involved in sorting of various transmembrane cargo proteins as well as transmembrane cargo receptors into nascent COPII vesicles (Miller et al., 2002; Miller et al., 2003; Mossessova et al., 2003; Peng et al., 1999). Membrane-bound Sar1-GTP/Sec23/24 complexes in turn recruit the outer COPII coat layer, the heterotetrameric Sec13/31 complex (Matsuoka et al., 1998). This is achieved by binding of an unstructured fragment within the proline-rich region of Sec31 to a binding site moulded by Sar1-GTP and Sec23, which explains the consecutive recruitment of the two coat layers (Bi et al., 2007). Polymerisation of Sec13/31 complexes leads to clustering of so-called pre-budding complexes composed of Sar1-GTP, Sec23/24 complexes and cargo, causing sculpting of the donor membrane into a vesicle bud and eventually leading to vesicle membrane scission.

All mammalian COPII coat subunits, except Sec13, are expressed in multiple isoforms. Four isoforms of the cargo binding subunit Sec24 were identified that can be grouped based on sequence homology into the two classes Sec24A/B and Sec24C/D (Pagano et al., 1999; Tang et al., 1999). Like their yeast homologues, (Miller et al., 2002; Miller et al., 2003; Peng et al., 1999; Shimoni et al., 2000) mammalian Sec24 isoforms function to expand the cargo repertoire of proteins that are actively sorted into COPII vesicles (Bonnon et al., 2010; Mancias and Goldberg, 2007, 2008).

Some cargo proteins can be taken up into nascent COPII transport vesicle by a passive mechanism termed bulk flow (Thor et al., 2009; Wieland et al., 1987). However, efficient ER export of a variety of transmembrane cargo proteins as well as transmembrane cargo receptors, in concert with their client proteins, relies on signal-mediated sorting into COPII vesicles. The short linear peptide motif with the consensus sequence DxE present in the cytoplasmic tail of VSVG was the first transport signal to be identified (Nishimura and Balch, 1997). A series of biochemical, genetic, and structural studies leads to the identification of various other short peptide motifs, as well as to their corresponding cargo binding sites on the Sec24 subunits. Sed5p harbours an YxxxNPF motif that promotes sorting of the yeast Qa-SNARE via the so-called A-site. The yeast Qc-SNARE Bet1p as well as VSVG are sorted into nascent COPII vesicles via an LxxLE or aforementioned DxE motif, respectively, by binding to the so-called B-Site (Mancias and Goldberg, 2008; Miller et al., 2003; Mossessova et al., 2003). Not all transport motifs identified today are conserved between yeast and humans. The mammalian homologoue of Sed5p, Syntaxin5, lacks an YxxxNPF motif, and instead binds via an IxM motif to a distinct cargo-binding site present on the mammalian isoforms Sec24C and Sec24D (Mancias and Goldberg, 2008). The binding sites for Sec22p, as well as its mammalian homologoue Sec22b, also referred to as C-site, recognize a conformational epitope rather than a short peptide motif, which is formed when the R-SNARE is in its closed conformation. In mammalian cells this signal is recognized by the isoforms Sec24A and Sec24B (Mancias and Goldberg, 2007; Mossessova et al., 2003). Another set of ER export signals are di-hydrophobic motifs found in the cytoplasmic tails of p24/Emp24/TMED family proteins (Belden and Barlowe, 2001; Dominguez et al., 1998; Fiedler et al., 1996; Nakamura et al., 1998), ERGIC53/Emp47p proteins (Kappeler et al., 1997; Nufer et al., 2002; Sato and Nakano, 2002), and the Erv41/46 complex (Otte and Barlowe, 2002). Controversial reports exist in the literature with regard to the binding site used for sorting of these proteins into nascent COPII-coated vesicles, and furthermore on which Sec24 isoform(s) this binding site is located. ERGIC53 is packed into nascent COPII vesicles by all Sec24 isoforms to a similar extent (Adolf et al., 2016; Mancias and Goldberg, 2007, 2008). On the other hand, transport of p24 proteins, which serve as adaptors for ER-export of GPI-anchored proteins, was reported to involve Sec24C and Sec24D in mammalian cells (Bonnon et al., 2010), as well as the corresponding isoform Lst1p in yeast (Castillon et al., 2011). A more recent study provides evidence at a structural level that p24 proteins are sorted by the so-called B-site (Ma et al., 2017).

Some cargo proteins require coincidence detection of sorting signals present on both, a classic transmembrane cargo adaptor and the client protein itself. The ATPase Yor1p, which harbours a DxE motif to bind the B-site for sorting into COPII vesicles, requires Erv14p, which makes contact to the D-site via the motif IFRTL (Pagant et al., 2015).

In order to obtain a more comprehensive view of the various interactions Sec 24 isoforms can undergo, we here characterize the proteomes of various types of COPII vesicles generated with recombinant coat proteins in semi-intact cells by SILAC based mass spectrometry.

## Results and Discussion

### Reconstitution and purification of COPII vesicles for SILAC-based proteomic profiling

In order to determine the proteome of mammalian COPII vesicles to get further insight into the role of the mammalian Sec24 isoforms in cargo packing, we have combined an *in vitro* COPII vesicle formation assay from semi-intact cells (SIC) (Adolf et al., 2013; Adolf et al., 2016; Mancias and Goldberg, 2007, 2008) with stable isotope labelling by amino acids in cell culture (SILAC)-based mass spectrometric analysis (Ong et al., 2002). For this purpose we modified the standard COPII vesicle budding assay by introducing density gradient centrifugation as further purification step to ensure reliable mass spectrometric analysis. In brief: 1) (SIC) were prepared by permeabilisation with digitonin and the cytosol was removed by repetitive washing in assay buffer, 2) SIC donor membranes were incubated with recombinant Sar1b, inner and outer coat complexes (Sec23/24 and Sec13/31), and GTP (or an non-hydrolysable GTP analogue where indicated), 3) newly formed vesicles were separated from the SIC donor membranes by medium speed centrifugation, and 4) finally COPII vesicles were depleted from excess free coat proteins and other contaminations by density gradient centrifugation (Fig.1A).

**Fig.1:**
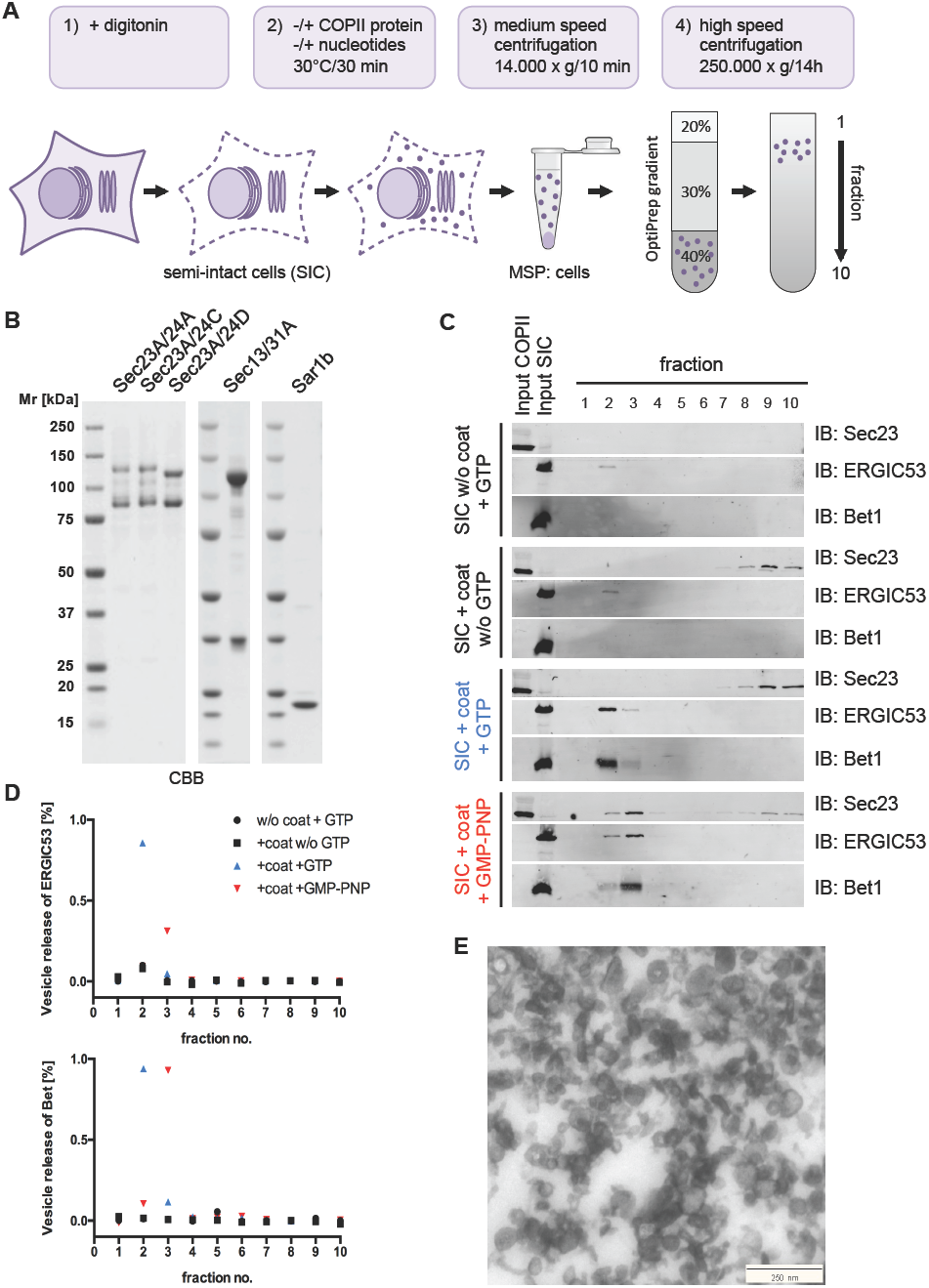
*In vitro* reconstitution of COPII vesicles from semi-intact cells for SILAC-based mass spectrometric analysis. A) Schematic diagram of the *in vitro* COPII vesicle reconstitution assay. Semi-intact cells (SIC) were incubated with Sar1b, Sec23/24, and Sec13/31 coat subcomplexes and guanine triphosphate (GTP). Newly formed vesicles were separated by medium speed centrifugation and subsequently further purified by floatation in an iodixanol gradient. After centrifugation COPII vesicle-containing fractions were harvested from top prior MS or WB analysis. B) Recombinant COPII coat proteins used in this study. Purified Sar1b (0.5 μg), the inner COPII coat subcomplexes His6-Sec23A/24A, His6-Sec23A/24C, or His6-Sec23A/24D (1 μg), and the outer coat subcomplex His6-Sec13/31A (1.5 μg) were separated by SDS polyacrylamide gel electrophoresis and stained with Coomassie brilliant blue (CBB). C) Density gradient enrichment of *in vitro* reconstituted COPII vesicles. COPII vesicles were reconstituted by incubation of SIC with Sar1b, Sec23A/24C, and Sec13/31A, in the presence of the indicated nucleotide (GTP or GMPPNP) and an ATP-regenerating system (ATPr). Newly formed vesicles were separated from donor membranes by medium speed centrifugation and subsequently floated in an iodixanol density gradient. Fractions from the gradient were harvested from top (1) to bottom (10) and analysed by Western blotting for the presence of the inner coat subcomplex subunit Sec23, and the two vesicle cargo proteins ERGIC53 and Bet1. The lanes labelled Input COPII and Input SIC represent 2.5% of starting material used in COPII vesicle reconstitutions. D) Quantification of COPII packing efficiency. Western blot signals of the two COPII cargo proteins ERGIC53 and Bet1 in the different fractions were quantified utilizing the Li-COR Image Studio software. E) Electron microscopic analysis of ultrathin sections of resin-embedded reconstituted COPII vesicles after gradient density purification. Prior resin embedding, fractions 2 and 3 were combined, diluted with assay buffer, and subsequently centrifuged to pellet COPII vesicles.

We initially intended to analyse the proteome of COPII vesicles generated with all four mammalian Sec24 isoforms. In addition to the complexes Sec23A/Sec24A and Sec23A/Sec24C, which have recently been described and used to investigate the Sec24 isoform-dependent sorting of ER-to-Golgi SNARE proteins (Adolf et al., 2013; Adolf et al., 2016), we cloned, expressed, and purified the complexes Sec23A/Sec24B and Sec23A/Sec24D (SI Fig.1). Non-stoichiometric amounts of free hexa-histidin-tagged Sec23 after affinity purification required further purification by gel filtration. Finally, we succeeded in obtaining satisfactory results with regard to yield and purity for the complexes Sec23A/Sec24A, Sec23A/Sec24C, and Sec23A/Sec24D (Fig.1B), and performed quantitative proteomic analysis of COPII vesicles reconstituted with any of these inner coat subcomplexes.

When *in vitro* generated vesicles were harvested by differential centrifugation and subsequently subjected to mass spectrometric analysis, unreliable results were obtained, probably due to the presence of the excess amounts of free coat proteins in the vesicle fractions. Therefore, we introduced a gradient density centrifugation step prior to pelleting of the vesicles. For this purpose vesicle-containing supernatants from COPII vesicle budding reactions were adjusted to 40% (w/v) iodixanol and overlaid with 30% (w/v) and 20% (w/v) iodixanol (OptiPrep) in assay buffer, and subsequently centrifuged for 14h. To determine the migration of COPII vesicles, ten fractions were collected form top (fraction 1) to the bottom (fraction 10) of the gradient and probed for the presence of the inner coat subunit Sec23A, and the two COPII vesicle cargo proteins ERGIC53 and Bet1 (Fig.1C). In control experiments SIC were incubated either without recombinant COPII coat proteins in the presence of GTP (Fig.1C, top panel) or with the recombinant COPII coat proteins (Sar1b, Sec23A/24C, Sec13/31A) in the absence of GTP (Fig.1C, top but one panel). These controls revealed some residual signal for ERGIC53 and Bet1 detected in fractions 2 and 3 (Fig.1C, upper two panels). These signals most likely stem from COPII and COPI vesicles formed by residual coat proteins that were not removed during preparation of the SIC, or from unspecific shearing of donor membranes. When SIC were incubated with recombinant COPII coat proteins plus either GTP (Fig.1C, bottom but one panel), or its nonhydrolysable analogue GMP-PNP (Fig.1C, bottom panel), clearly elevated signals for the two Sec24C cargo proteins ERGIC53 and Bet1 were observed in fractions 2 and 3 of the gradient. To prevent COPII vesicle coat disassembly after vesicle membrane scission we substituted GTP with GMPPNP. Consistently, the inner COPII coat subunit Sec23A was also detected in these fractions 2 and 3, when the non-hydrolysable GTP analogue was utilised (Fig.1C, bottom panel).

Furthermore, and in line with the higher buoyant density of COPII-coated vesicles, the majority of the vesicles were recovered at slightly higher density (fraction 3) when GMP-PNP was used, whereas *in vitro* reconstitutions with GTP yield uncoated COPII vesicles, mainly recovered from fractions of lower density (fraction 2) (Fig.1C, compare lanes 2 and 3). The relative quantity of the COPII vesicle cargo proteins ERGIC53 and Bet1 along the different gradients is shown in Fig.1D. To further characterise the COPII vesicle fraction reconstituted from SIC and enriched by flotation, combined gradient fractions 2 and 3 were resin-embedded and examined by transmission electron microscopy. Predominantly, coated vesicular structures of a size range between 60 and 100 nm were observed (Fig.1E).

### SILAC-based quantitative mass spectrometric analysis of COPII vesicles reconstituted with distinct Sec24 isoforms

For quantitative mass spectrometric analysis of COPII vesicles we combined *in vitro* COPII vesicle reconstitution as described above with stable isotope labelling in cell culture (SILAC). The general workflow for SILAC-based quantitative proteomics analysis of COPII vesicles from semi-intact cells (SIC) is depicted in Fig.2A: 1) cells were cultured in either ‘light’ medium with natural amino acids (Lys-0/Arg-0) or in ‘heavy’ medium with C13/N15-labelled amino acids (Lys-8/Arg-10), 2) SIC were prepared from ‘light’ and ‘heavy’ cells for *in vitro* reconstitution of COPII vesicles, 3) vesicles were isolated as described above, and 4) corresponding vesicle samples were mixed and analysed by MS (e.g. vesicle fractions generated with isoform X from ‘light’ cells were mixed with vesicle fractions generated with isoform Y from ‘heavy’ cells). To compensate for any effect metabolic labelling may have on the uptake of cargo proteins into COPII vesicles, all assays were conducted as label-switch experiment.

For proteome analysis, COPII vesicles were reconstituted with either Sec23A/24A, Sec23A/24C, or Sec23A/24D, and mixed with the corresponding fraction obtained after incubation of SIC without recombinant coat proteins in the presence of GTP and an ATP regenerating system (mock reaction). To assess the extent of vesicle reconstitution, a small fraction of each sample was probed by Western blotting prior to mixing for the presence of the ER resident protein Calnexin (negative control), and of known COPII marker proteins (positive control). As indicated in Fig.2B, vesicles reconstituted with Sec23A/24C from ‘light’ cells were mixed with the mock vesicle reconstitution reaction from ‘heavy’ cells and vice versa (Fig.2B). In general, no differences were observed with regard to the yield and/or specificity of packaging of cargo or non-cargo proteins into COPII vesicles, irrespectively whether unlabelled or metabolically labelled cells were used for vesicle reconstitution (Fig.2B). We first defined the mammalian COPII vesicle core proteome by combining COPII vesicle fractions reconstituted with the inner coat subcomplexes Sec23A/24A (Fig.2C and F), Sec23A/24C (Fig.2D and G), or Sec23A/24D (Fig.2E and H) with mock reactions. All reactions were carried out as label-switch experiments as described above and the relative abundance of proteins was quantified with MaxQuant (SI Table 1). Proteins with a log2 SILAC ratio clustering around zero are similarly abundant in both the vesicle reconstitution reaction and the mock reaction and hence are regarded as contaminants. In reconstitutions with Sec23A/24A 51, with Sec23A/24C 76, and with Sec23A/24D 69 proteins were quantified with an average log2 SILAC ratio >1. Proteins quantified in corresponding vesicles from ‘heavy’ or ‘light’ labelled cells with an log2 SILAC ratios > 0.5 are depicted in Fig.2 C-E. Corresponding scatter plots of all quantified proteins in the different SILAC label-switch experiments are shown in SI Fig.2. Proteins known to function in the early secretory pathway are highlighted in red and labelled with their gene names. In Fig 2 the top 30 proteins enriched in COPII vesicle fractions reconstituted with Sec23A/24A (F), Sec23A/24C (G), and Sec23A/24D (H) are ranked according to their average log2 SILAC ratio.

**Fig.2:**
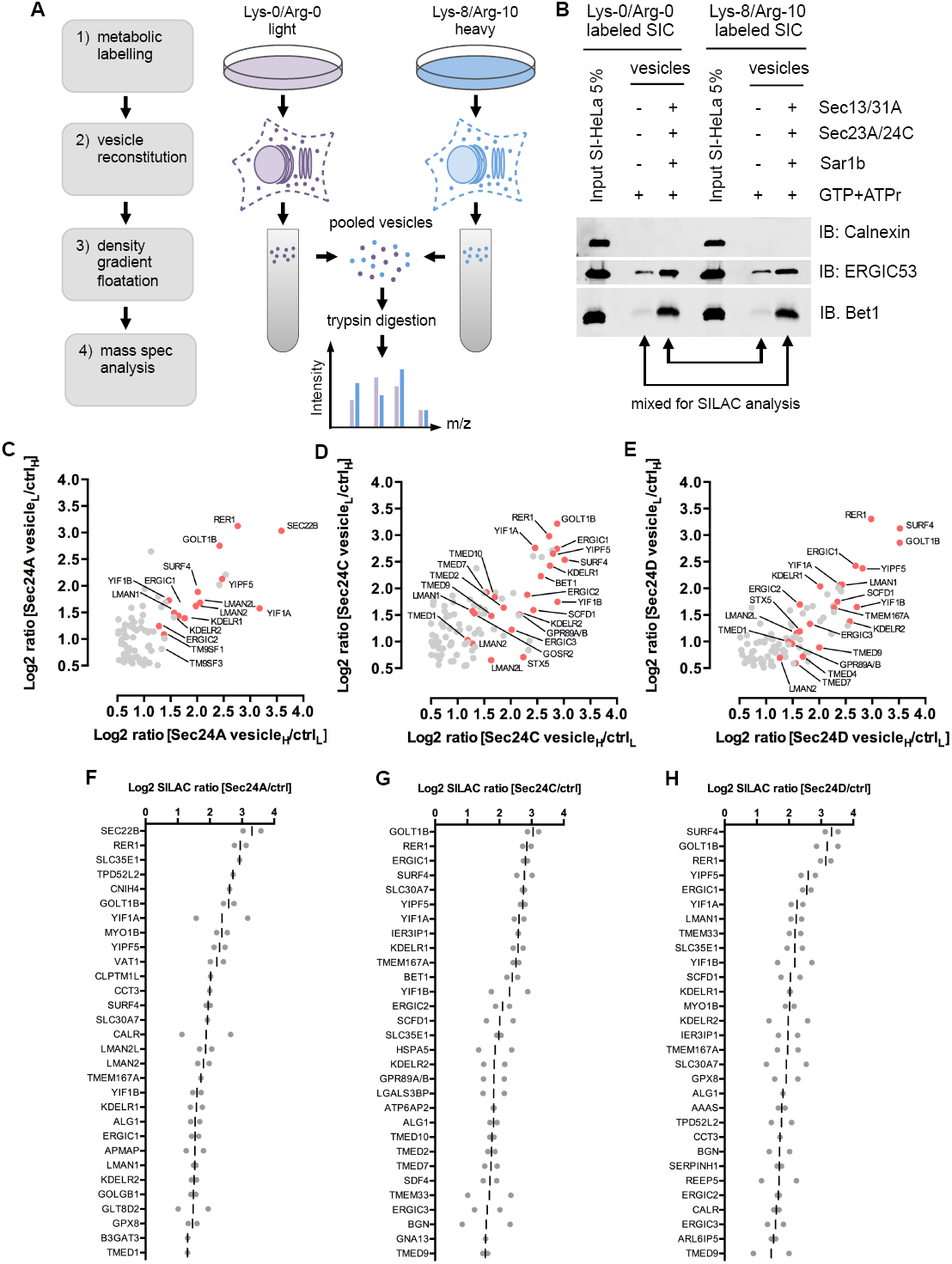
SILAC-based proteomic profiling of isotopic COPII vesicles. A) Schematic diagram of the workflow for SILAC-based mass spectrometric analysis of COPII vesicles. HeLa cells were metabolically labelled with ‘light’ (Lys-0/Arg-0) or ‘heavy’ (Lys-8/Arg-10) amino acid isotypes (1). COPII vesicles were reconstituted with SIC prepared form ‘light’ or ‘heavy’ labelled cells (2), and subsequently enriched by floatation, and pooled with samples from a corresponding control budding reaction (SIC+GTP) (3), prior to mass spectrometric quantification (4). B) Western blot analysis of label-switch experiment. SIC from ‘light’ and ‘heavy’ labelled cell were incubated with recombinant COPII coat proteins as indicated and subsequently analysed by Western blotting for the presence of the ER-resident (non-cargo) marker Calnexin and the two COPII vesicle cargo proteins ERGIC53 and Bet1. C-H) Proteomics of *in vitro* reconstituted isotopic COPII vesicles. Scatter plot depicting proteins quantified with an log2 SILAC ratio > 0.5 in SILAC label-switch experiments of COPII vesicles reconstitutions from ‘light’ or ‘heavy’ labelled SIC with the isoforms Sec23A/24A (C), Sec23A/24C (D), or Sec23A/24D (E). COPII cargo proteins and proteins which cycle in the early secretory pathway are highlighted in red and labelled with their gene name. Log2 SILAC ratios of label-switch experiments of COPII vesicle reconstitutions with the isoforms Sec23A/24A (F), Sec23A/24C (G), or Sec23A/24D (H) were calculated and proteins were ranked from highest to lowest ratio. Horizontal lines indicate the mean ratio.

All proteins identified with an elevated log2 SILAC ratio are known to function in the early secretory pathway or have been identified as ER-Golgi trafficking proteins. Among these are 1) Surf4, the mammalian homologoue of the ER-export adaptor Erv29p; 2) Rer1, known to function as adaptor protein for retrograde transport of escaped ER transmembrane proteins; 3) Golt1B, an ER-Golgi cycling protein with a proposed function in COPII vesicle tethering/fusion; 4) the ER-Golgi-SNARE proteins Sec22b, Syntaxin5, GS27, and Bet1; 5) proteins of the ERGIC53/LMAN1 family; 6) proteins of the ERGIC1/2/3 family; 7) proteins of the Yip/Yif family; 8) proteins of the p24/TMED family; and 9) the KDEL receptor family (Fig.2 C-E).

Consistent with earlier reports, (Adolf et al., 2016; Mancias and Goldberg, 2007, 2008), significantly different log2 SILAC ratios were obtained for the ER-to-Golgi SNARE proteins. Sec22b scored the highest SILAC ratio in reconstitutions with the isoform Sec24A but showed only slight or no enrichment in reconstitutions carried out with either Sec24C or Sec24D. On the other hand, all three Q-SNAREs displayed a higher log2 SILAC ratio when reconstitutions were carried out with Sec24C or Sec24D compared to Sec24A. Utilizing *in vitro* reconstitutions in combination with quantitative Western blotting, several reports have shown that ERGIC53 is packaged into mammalian COPII vesicles in a Sec24-isoform independent manner (Adolf et al., 2016; Mancias and Goldberg, 2007, 2008). Consistently, in our proteomic data ERGIC53/LMAN1 displayed similar average log2 SILAC ratios in vesicles generated with Sec24A (1.53) and Sec24C (1.43), and a slightly higher log2 SILAC ratio when experiments were conducted with Sec24D (2.23) (Fig.2 C-E). Further confirming previous reports, members of the p24/TMED family, which were reported to be transported selectively by the isoforms Sec24C/D, (Bonnon et al., 2010) were reproducibly found enriched in vesicles generated with the isoform Sec24C and Sec24D, but not Sec24A. Moreover, while no difference in enrichment was observed of the proteins Gollt1B, of members of the YIP/YIF or KDEL-receptor family, the proteins ERGIC1 and Surf4 showed reproducibly higher log2 SILAC rations in vesicle reconstitutions with Sec24C and Sec24D compared to Sec24A (Fig.2 C-H). To further investigate which cargo proteins are transported in a Sec24 isoform-specific manner, we directly compared vesicle fractions generated with Sec23A/24A or with Sec23A/Sec24C (Fig.3A/B). Based on the log2 SILAC ratios of two independent label-switch experiments (n=4) p-values were calculated and plotted against the mean log2 SILAC ratios (Fig.3B and SI Table S2). Consistent with previous reports and our own observations, (Adolf et al., 2016; Mancias and Goldberg, 2007, 2008) ERGIC53/LMAN1 is packaged into isotypic Sec24A and Sec24C COPII vesicles at approximately the same level (Fig.3B). The only two proteins enriched in Sec24A vesicle fractions are the ER-to-Golgi R-SNARE Sec22b (Fig.3B, highlighted in green), and CNIH4 (Fig.3B, highlighted in red). Sec22b is a known Sec24A/B cargo protein (Mancias and Goldberg, 2007), which is sorted into COPII vesicles by binding to the C-site of the Sec24 subunit (Miller et al., 2003; Mossessova et al., 2003). It is of note that CNIH4 was also one of the top hits (log2 SILAC ratio=2.6 vesicle_H_/ctr_L_), when the core proteome of vesicles reconstituted with the mammalian isoform Sec24A was determined (Fig.2F). CNIH4 and its yeast homologue Erv14p are trans-membrane cargo adaptor proteins for GPCRs or Axl2p, respectively (Powers and Barlowe, 1998, 2002; Sauvageau et al., 2014). More recently, Erv14p was reported to be involved in the sorting of various transmembrane proteins by coincidence detection, where sorting signals present on the cytoplasmic tails of Erv14p together with a cargo protein form a proper bivalent motif (Pagant et al., 2015). This study also revealed a novel cargo-binding site on Sec24p, designated D-site, which is utilized by Erv14p. The critical residues within this site (Sec24p: S^491^, F^576^, and R^578^), involved in binding of the IFRTL sorting motif of Erv14p, are conserved in the mammalian isoforms Sec24A/B, but not in Sec24C/D (Fig.3C). Taken together, these data suggest that CNIH4 is a novel Sec24A/B-dependent cargo adaptor.

**Fig.3:**
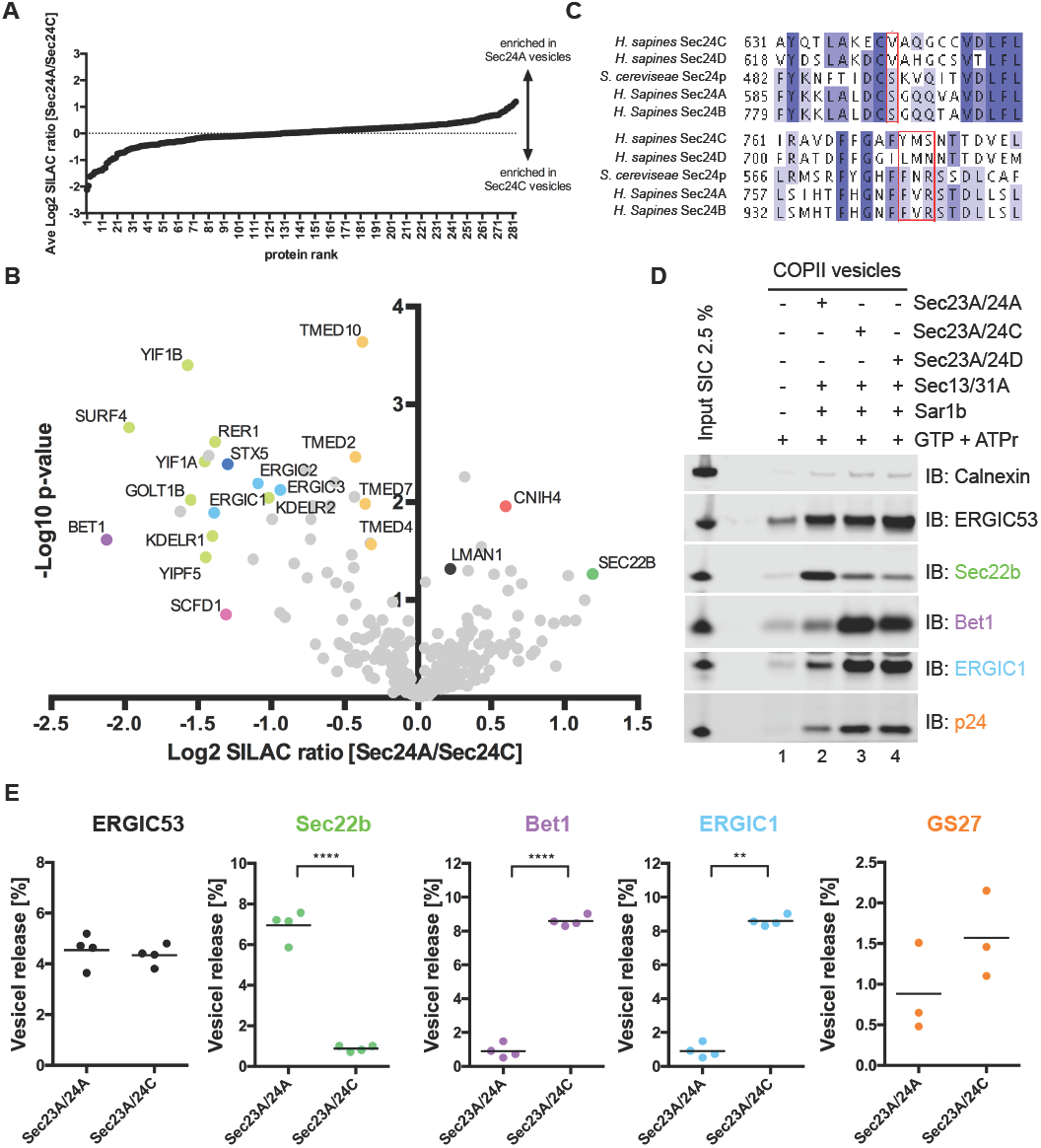
Identification of novel Sec24-isoform specific cargo proteins. A-B) Direct proteomic comparison of isotypic COPII vesicles reconstituted with inner coat subcomplexes containing Sec24A or Sec24C. A) Log2 SILAC ratios of label-switch experiments of vesicle reconstitutions with Sec23A/24A and Sec23A/24C were calculated and quantified cargo proteins were ranked from lowest (enriched in Sec24C vesicles) to highest (enriched in Sec24A vesicles) ratio. B) Volcano plot of two independent label-switch experiments (n=4) of a direct comparison of Sec24A vesicles with Sec24C vesicles. Log2 SILAC ratios (Sec24A/Sec24C) were calculated and plotted against their respective p-values (-log10). Proteins of interest are color-coded and labelled with their gene names. Sec24C-specific cargo proteins are colored in magenta, and Sec24A-specific cargo proteins are colored in red. ERGIC53/LMAN1, transported by all isoforms, is colored in light green. Putative novel Sec24C-dependent cargo proteins are colored in blue and putative novel Sec24A-dependent cargo proteins are colored in orange. C) Critical residues for cargo binding within yeast Sec24p are conserved in human Sec24A/B. Sequences of human Sec24A, B, C, and D, as well as yeast Sec24p were aligned with clustal omega (EMBL-EBI) using the default settings. Depicted are protein sequence segments showing that the critical residues within the previously identified D-site (Sec24p: S^491^, F^576^, and R^578^, red box) are conserved in human Sec24A and Sec24B, but not in human Sec24C and Sec24D. D-E) Validation of ERGIC1 as a novel Sec24C-dependent cargo protein. D) COPII-isoform dependent cargo packing efficiency of selected cargo proteins. COPII vesicles were reconstituted *in vitro* by incubation of semi-intact cells (SIC) with the Sar1b, the outer coat complex Sec13/31A and one of the inner coat complexes Sec23A/24A, Sec23A/24C, in the presence of GTP and an ATP-regenerating system (ATPr) as indicated and subsequently vesicles were separated from donor membranes by differential centrifugation. The vesicle fractions (40%) and input of semi-intact cells (SIC, 2.5%) used for reconstitution were analysed by Western blotting for the presence of the non-vesicle membrane marker Calnexin, the Sec24 isoform independent cargo protein ERGIC53, the ER-to-Golgi R-SNARE Sec22b (Sec24A/B-dependent), the ER-to-Golgi Qc-SNARE Bet1 (Sec24C/D-dependent), p24/TMED2, and ERGIC1. E) Quantification of COPII-isoform dependent cargo packing efficiency. The amount of the non-vesicle membrane marker Calnexin and the indicated COPII cargo proteins in vesicle fractions was quantified utilizing the Li-COR Image Studio software (means ± SEM, n=3-4).

Several proteins that we initially identified as core proteome of mammalian COPII vesicles (Fig.2) are enrichment in Sec24C over Sec24A vesicles (Fig.3A and B). To compensate potential mixing errors when combining the different samples, the MaxQuant integrated normalization method was applied. Furthermore, based on the following observations it can be ruled out that simply different amounts of Sec24C and Sec24A vesicle fractions were mixed prior mass spectrometric analysis: 1) the well-established Sec24A-dependent cargo protein Sec22b (highlighted in green) was found enriched in Sec24A vesicles, and 2) the isoform-unspecific cargo protein ERGIC53/LMAN1 (highlighted in black) was equally abundant in Sec24A and Sec24C vesicles.

Among the known Sec24C-specific cargo proteins Syntaxin5 and Bet, the cytosolic SM protein Sec1 family domain-containing protein 1 (SCFD1) (Fig.3B, highlighted in pink) was also enriched in Sec24C vesicles.

To further validate the results from the SILAC-based approach, we selected the ER-Golgi cycling proteins ERGIC1-3, because results from both experimental setups 1) comparison of Sec24-isoform specific vesicles with a mock reaction (Fig.2), and 2) comparison of Sec24C with Sec24A vesicles (Fig.3) implied that the mammalian homologues of Erv41p/Erv46p are novel Sec24C/D specific cargo proteins. All three proteins were reproducibly enriched in Sec24C over Sec24A vesicles (Fig.3B, highlighted in light blue). Consistently, for all three proteins higher log2 SILAC ratios where obtained when Sec24 isoform-specific vesicle fractions where compared to mock reactions (Fig.2 C-H). The combined average log2 SILAC ratio for ERGIC1, 2 and 3 in these experiments was for Sec24A vesicles 1.187 (SD 0.360), Sec24C vesicles 2.176 (SD 0.555), and for Sec24D vesicles 1.935 (SD 0.469) (Fig.2 and SI Table 1). For further validation, we conducted COPII vesicle reconstitutions in the presence of Sec23A/24A, Sec23A/24C, or Sec23A/24D and analysed vesicle fractions by Western blotting (Fig.3D). The non-cargo protein Calnexin served as negative control and the two known Sec24 isoform-specific cargo proteins Sec22b (Sec24A/B cargo) and Bet1 (Sec24C/D cargo) served as positive controls for specificity of cargo packing. When COPII vesicle fractions were analysed for the presence of ERGIC1, a clear preference for the isoforms Sec24C/D was observed (Fig.3D, compare lanes 2 with lanes 3 and 4).

Furthermore, we found p24/TMED family proteins to be slightly enriched in Sec24C/D COPII vesicles (Fig.2C-H and Fig.3B). Consistently, when COPII vesicles generated with different Sec24 isoforms were analysed by Western blotting, we observed more p24/TMED2 in COPII vesicles reconstituted with Sec24C and Sec24D compared to COPII vesicles reconstituted with Sec24A (Fig.3C, compare lanes 2 with lanes 3 and 4). These results are in agreement with an earlier report of Sec24C/D isoform-dependent sorting of p24/TMED proteins by the mammalian COPII coat (Bonnon et al., 2010). Independent replicates of COPII vesicle reconstitutions with either Sec24A or Sec24C are quantified in Fig.3E. Whereas Surf4, Golt1B, Rer1, Yif1A, Yif1B, Yipf5 and KDEL receptor 1 and 2 all were enriched in Sec24C over Sec24A vesicles, these proteins where also quantified with elevated log2 SILAC ratios in Sec24A vesicles measured against a mock reaction (Fig.2C and F), and thus require additional analysis before these proteins can be confirmed as additional novel Sec24C-dependent cargo proteins.

### Identification of novel client proteins for the Sec24C/D IxM cargo binding site

In order to test whether our experimental setup can also be used to investigate which cargo proteins are sorted into nascent COPII vesicles by a specific cargo-binding site we made use of the previously characterised IxM binding site located on Sec24C/D (Mancias and Goldberg, 2008). Mutation of this site results in defects in packing of the three Q-SNAREs Syntaxin5, GS27, and Bet1 (Adolf et al., 2016; Mancias and Goldberg, 2008). Previously, we have reported that these three ER-to-Golgi Q-SNAREs are sorted into COPII vesicles as a preassembled complex via the IxM motif residing in the linker region of Syntaxin5 between its N-terminal H_abc_-domain and the SNARE motif (Adolf et al., 2016). We compared vesicle fractions generated with Sec24C wild type and the Sec24C^LIL895AAA^, a synthetic mutant (Mancias and Goldberg, 2008) with a compromised IxM cargo-binding site (Fig.4A). Seven proteins were reproducibly depleted in COPII vesicles formed by the mutant coat protein (Fig.4B and C, and SI Table 2). Consistent with our previous report, all three ER-to-Golgi Q-SNAREs (Syntaxin5, GS27, and Bet1, Fig.4B) were among the proteins depleted in Sec24C^LIL895AAA^-vesicles. Notably, alongside the three SNARE proteins, the SM protein SCFD1 (Fig.4B), homologue of the *Saccharomyces cerevisiae* SM protein Sly1p, was also considerably decreased in Sec24C^LIL895AAA^ vesicles (Fig.4B and C). This observation is in line with our results above. The SM protein was among the top 30 hits in Sec24C and Sec24D vesicles, but not in Sec24A vesicles (Fig.2F-H), and was furthermore found to be enriched in Sec24C vesicles when directly compared with Sec24A vesicles (Fig.3B). Taken together, this suggests that the SM protein SCFD1 binds the ER-to-Golgi Q-SNARE complex prior to its packing into COPII vesicles.

**Fig.4:**
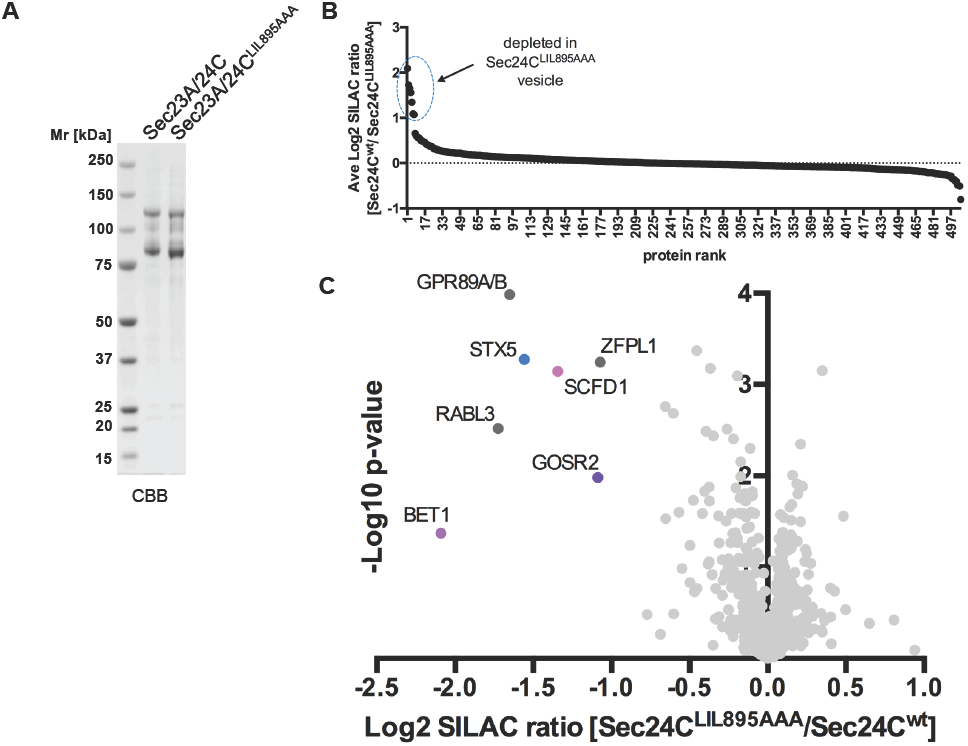
Identification of novel clients of the Sec24C/D IxM cargo-binding site. A) Sec23A/24C wild type and Sec23A/24C^LIL895AA^ inner COPII coat subcomplexes. Recombinant inner coat subcomplexes (1 μg) were separated by SDS polyacrylamide gel electrophoresis and stained with Coomassie brilliant blue (CBB). B-C) Direct proteomic comparison of COPII vesicles reconstituted with inner coat subcomplexes containing Sec24C^LIL895AAA^ or Sec24C wild type proteins. A) Log2 SILAC ratios of label-switch experiments of vesicle reconstitutions with Sec23A/24C wild type and Sec23A/24C^LIL895AAA^ were calculated and quantified cargo proteins were ranked from highest to lowest ratio. In total seven proteins showed a mean log2 SILAC ratio (Sec24C wt/Sec24mutant) above 1. C) Volcano plot of three independent label-switch experiments (n=6) of a direct comparison of COPII vesicles generated with Sec24C wild type vesicles or Sec24C^LIL895AAA^. Log2 SILAC ratios (Sec24Cwt/Sec24C^LIL895AAA^) were calculated and plotted against their respective p-values (-log10). Proteins of interest are color-coded and labelled with their gene names. ER-to-Golgi QSNARE proteins and the SM protein SCFD1 are colored in red. Other proteins depleted in vesicles generated with the Sec24C mutant protein are colored in dark grey.

Three more proteins (Fig.4C, highlighted in dark grey) are depleted from vesicles reconstituted with the Sec24^LIL895AAA^, the Golgi pH regulator A/B (GPR89 A/B), the Zinc finger protein-like 1 (ZFPL1), and Rab-like protein 3 (RabL3).

### Biochemical characterization of Sec24D mutants implicated in the development of osteogenesis imperfecta

Mutations within proteins involved in COPII vesicle biogenesis were repeatedly identified as the origin of various severe medical conditions (Boyadjiev et al., 2006; Jones et al., 2003; Yehia et al., 2015). Most recently, mutations in Sec24D were reported to be implicated in the development of a syndromic form of osteogenesis imperfecta (OI), an affliction usually coincidental with mutations in genes of type I collagen. These mutations were observed in close vicinity to the known IxM cargo-binding site mentioned above (Garbes et al., 2015). We therefore decided to investigate effects *in vitro* of the two single point mutations QE978P and S1015F reported in Sec24D. To this end, Sec23/Sec24D complexes carrying the respective mutations were expressed and purified (Fig.5A). The positions of both mutations are shifted by one amino acid (Q979P and S1016F), because isoform 2 of Sec24D with an additional alanine in position 224 was used.

**Fig.5:**
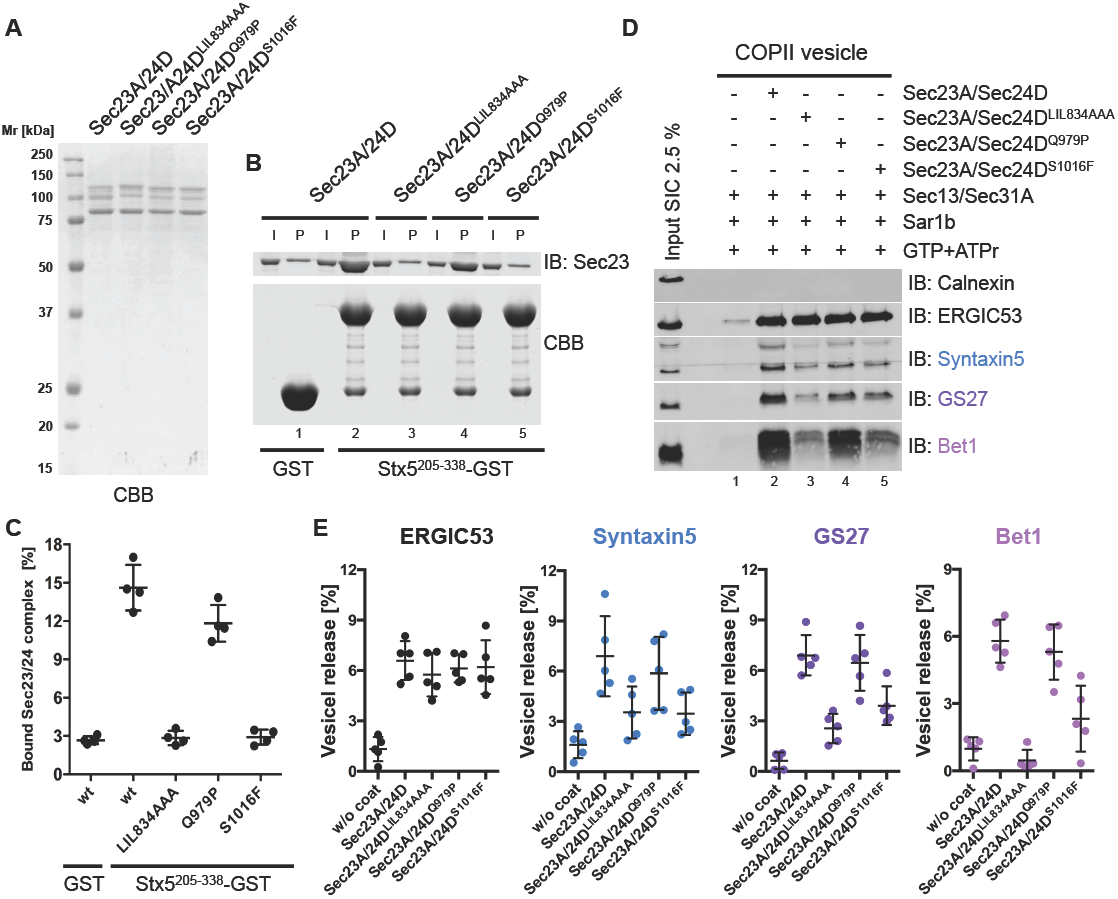
Biochemical analysis of Sec24D mutant proteins implicated in the development of a syndromic form of osteogenesis imperfecta. A) Sec23A/24D wild type, Sec23A/24D^LIL895AA^, Sec23A/Sec24D^Q979P^, and Sec23A/Sec24D^S1016F^ inner coat subcomplexes. Recombinant inner coat subcomplexes (1μg) were separated by SDS polyacrylamide gel electrophoresis and stained with Coomassie brilliant blue (CBB). B) The Sec24D^S1016F^ has severely compromised ability to directly bind an IxM sorting signal containing Syntaxin5 fusion protein. Direct interaction of the ER-to-Golgi Qa-SNARE fusion proteins of Syntaxin5 wt and different Sec23A/24D mutants was probed by GST pull down assay. Sec23A/24D complexes bound to GST or Syntaxin5^205-338^-GST (P) as well as inputs (2%) of inner coat subcomplexes were separated by SDS polyacrylamide gel electrophoresis. The lower part of the polyacrylamide gel containing GST and Syntaxin5^205-338-^GST fusion protein was stained with Coomassie brilliant blue (CBB), and the upper part was analysed by Western blotting for the presence of Sec23A. C) Quantification Syntaxin5^205-338^-GST pull-down experiments. Amount of Sec23A/24D coat subcomplexes bound to GST or Syntaxin5^205-338^-GST was quantified utilizing the Li-COR Image Studio software (means ± SEM, n=4). D) Cargo packing efficiency of Sec24D mutant proteins implicated in the development of osteogenesis imperfect. COPII vesicles were reconstituted *in vitro* by incubation of semi-intact cells (SIC) with the Sar1b, the outer coat complex Sec13/31A and Sec23A/24D inner coat complexes containing wild type or the indicated Sec24D-mutant protein, in the presence of GTP and an ATP-regenerating system (ATPr). COPII vesicles were separated from donor membranes by differential centrifugation. The whole vesicle fractions and input of semi-intact cells (SIC, 2.5%) used for reconstitution were analysed by Western blotting for the presence of the non-vesicle membrane marker Calnexin, the Sec24 isoform independent cargo protein ERGIC53, and the ER-to-Golgi Q-SNAREs Syntaxin5, GS27, and Bet1. E) Quantification Sec24D mutant ER-to-Golgi Q-SNARE packing efficiency. The amount of the non-vesicle membrane marker Calnexin, ERGIC53, as well as the amount of the three ER-to-Golgi Q-SNARE proteins Syntaxin5, GS27 and Bet1 in vesicle fractions generated with the indicated Sec24D proteins was quantified utilizing the Li-COR Image Studio software (means ± SEM, n=5).

Taking advantage of a previously established pull down assay using Syntaxin5^205-328^-GST as bait, we tested the capability of the mutant coats to bind to the critical IxM SNARE-sorting motif in Syntaxin5 (Adolf et al., 2016; Mancias and Goldberg, 2008). Wild type Sec23/Sec24D complex was efficiently pulled down (Fig.5 B and C). A comparable interaction was observed for Sec23/Sec24D^Q979P^ (Fig.5 B and C). Binding of the Syntaxin5 fragment to either the synthetic mutant Sec23/Sec24D^LIL834AAA^ or a complex carrying the serine to phenylalanine (Sec23/Sec24D^S1016F^) point mutation, however, was completely abrogated as it dropped to the level of the GST control (Fig.5B and C).

In order to investigate functionally the impaired SNARE interaction of Sec23/Sec24D^S1016F^, variant and wild type coat proteins were compared by Western Blotting of COPII vesicles reconstituted *in vitro* (Fig.5 D). The non-cargo marker Calnexin was absent from all vesicle fractions, independent of the inner coat complex used, and the COPII cargo ERGIC53 was found with similar abundance in vesicles formed by either wild type or mutant Sec23/Sec24D complexes (Figs.5 D and E). In line with the pull-down results (Fig5, B and C), sorting of Syntaxin5 into COPII vesicles was largely impaired by the point mutation S1016F (Fig.5 D and E). This SNARE-packaging defect affects not only Syntaxin5, but also the other two Q-SNAREs Bet1 and GS27 (Fig.5D and E), and is similar to the loss of SNAREs observed for the synthetic mutant Sec23/Sec24D^LIL834AAA^. Again, these observations are consistent with the notion that the ER-Golgi Q-SNAREs are recruited into COPII vesicles as a complex (Adolf et al., 2016).

### Conclusions

We report a comprehensive analysis of the proteomes of HeLa cell-derived COPII vesicles by combining *in vitro* reconstitution with recombinant coat proteins and SILAC-based quantitative mass spectrometry. We have defined a set of 50-70 proteins as the core constituents of mammalian COPII vesicles. All of these proteins were previously identified as residing in the early secretory pathway. Comparison of the proteomes of vesicles reconstituted with defined isoforms of Sec24 allowed us to validate several cargo proteins to be transported in a Sec24 isoform-specific manner. Furthermore, we have identified and validated ERGIC1 as a novel Sec24C/D isoform-specific cargo protein, and provide evidence that CNIH4 is sorted by Sec24A.

Analysis of Sec24C/D mutant proteins with a compromised IxM binding site revealed that the SM protein SCFD1 binds to ER-localized Syntaxin5, where it is possibly involved in stabilizing the open conformation of the Qa-SNARE, prerequisite for Q-SNARE complex assembly and sorting into COPII-coated vesicles. Furthermore, analysis of Sec24D mutants implicated in the development of a syndromic form of osteogenesis imperfecta (Garbes et al., 2015) showed sorting defects for the three ER-to-Golgi Q-SNAREs Syntaxin5, GS27, and Bet1, similar to Sec24C/D mutants with compromised IxM cargo-binding site.

The methodology applied here can be used to tackle a variety of other cell biological questions. More recently, the ER-Golgi intermediate compartment (ERGIC) as donor membrane source as well as COPII coat components have been implicated to play a direct role in autophagosome formation (Ge et al., 2013; Ge et al., 2014; Lemus et al., 2016). It would be of particular interest to analyse the proteome of ERGIC-derived COPII vesicles to obtain insight into their compositions and thus to identify proteins involved in their formation. More recently we have utilized semi-intact cells to reconstitute COPI vesicles *in vitro* (Adolf et al., 2013; Adolf and Wieland, 2013). The assay described here was used to dissect the proteome of COPI vesicles and to further investigate potential distinct roles of the different COPI isoforms in cargo sorting (Rhiel et al.,bioRxiv 254052). The combination of *in vitro* reconstitution of vesicles from semi-intact cells together with SILAC-based quantitative proteomic analysis could be further extended to other vesicle types, e.g. AP1/GGA-dependent Clathrin-coated vesicles operating at the trans-Golgi network (TGN)-endosome interface, or AP4-coated vesicles involved in the sorting of cargo at the TGN.

## Materials and Methods

### Antibodies

The following antibodies were used: Calnexin (ab75801) from Abcam (Cambridge, MA); Sec23 (E19, sc-12107), ERGIC53 (C-6, SC-365158), Syntaxin5 (B-8, SC-365124), Gs37 (25, SC-135932), Sec22b (29-F7, SC-101267), and Bet1 (17, SC-136390) all from Santa Cruz Biotechnology (Dallas, TX); ERGIC1 (16108-1-AP) from Proteintech (Manchester, UK); p24 (Gommel et al., 1999). The following secondary antibodies were used: goat anti-rabbit Alexa Fluor 680 and goat anti-mouse Alexa Fluor 680, both from Invitrogen (Carlsbad, CA).

### Plasmids

The pFBDM transfer vector (Berger et al., 2004) as well as the plasmids pFBDM-His6-Sec23A, pFBDM-His6-Sec23A/Sec24A, pFBDM-His6-Sec23A/Sec24C, pFBDM-His6-Sec23A/Sec24C^LIL895AAA^, and pFBDM-His6-Sec13/Sec31A (Adolf et al., 2013; Adolf et al., 2016) have been described previously. The cDNAs of human Sec24B and human Sec24D were obtained as I.M.A.G.E. clones from Source BioScience (Nottingham, UK). For generation of the pFBDM-His6-Sec23A/Sec24B construct, the Sec24B cDNA was amplified by PCR with the primer 5´-AAA AAG AGC TCA TGT CGG CCC CCG CCG GGT C-3´ and 5´-AAA AAG CGG CCG CTC ACT TAC AAA TCT GCT GCT-3´, digested with SacI and NotI, and inserted into the respective restriction sites of pFBDM-His6-Sec23A. For generation of the pFBDM-His6-Sec23A/Sec24D construct, the Sec24D cDNA was amplified by PCR with the primer 5´-AAA AAG CGG CCG CAT GAG TCA ACA AGG TTA CGT G-3´ and 5´-AAA AAT CTA GAT TAA TTA AGC AGC TGA CAG AT-3´, digested with NotI and XbaI, and inserted into the respective restriction sites of pFBDM-His6-Sec23A. The pFBDM-His6-Sec23A/Sec24D^LIL834AAA^, pFBDM-His6-Sec23A/Sec24D^Q979P^, and pFBDM-His6-Sec23A/Sec24D^S1016F^ constructs were generated by site-directed mutagenesis.

### Protein expression and purification

The COPII coat subcomplexes were expressed in Sf9 insect cells and purified as described previously (Adolf et al., 2013; Adolf et al., 2016). Glutathione-S-transferase tagged hamster Sar1b was expressed in *E. coli* BL21 (DE3) pLysS (Thermo Fisher Scientific, Waltham, MA), and purified as described previously (Kim et al., 2005). The Syntaxin5^205-328^-Glutathion-S-transferase fusion protein as well as Glutathion-S-transferase were expressed in *E. coli* BL21 (DE3) (Thermo Fisher Scientific, Waltham, MA), and purified as described previously (Adolf et al., 2016).

### COPII vesicle formation from semi-intact cells and density gradient isolation

For standard Western blotting analysis of COPII vesicles HeLa cells ACC57 (DSMZ, Braunschweig, Germany) were cultivated at 37°C with 5% CO_2_ in alpha-MEM (Sigma-Aldrich, St. Louis, MO) with 10% fetal bovine serum. Preparation of semi-intact HeLa cells (Mancias and Goldberg, 2007) and the COPII budding assay was carried out as described previously (Adolf et al., 2013). For COPII vesicles formation and subsequent floatation 200 μg semi-intact HeLa cells were incubated for 30 min at 30°C in 200 μl assay buffer [25 mM HEPES pH 7.2 (KOH), 150 mM KOAc, 2 mM MgOAc] in the presence of an ATP-regenerating system (ATPr) [40 mM creatine phosphate (Roche, Penzberg, Germany), 0.2 mg/ml creatine phosphokinase (Sigma-Aldrich, St. Louis, MO), 1 mM ATP (Sigma-Aldrich, St. Louis, MO)], and 0.5 mM GTP (Sigma-Aldrich, St. Louis, MO) or 0.1 mM GMP-PNP (Sigma-Aldrich, St. Louis, MO), in the presence of 4 μg Sar1b, 8 μg Sec23/24 inner coat complex (Sec24A/24A, Sec23A/24C, or Sec23A/24D) and 10 μg Sec13/31A outer coat complex. After incubation, newly formed vesicles were separated from donor semi-intact cells by a medium speed centrifugation at 14.000 × g for 10 min at 4°C. The vesicle-containing supernatant was harvested, adjusted to 40% (w/v) iodixanol (OptiPrep^TM^, Sigma-Aldrich, St. Louis, MO) in a total volume of 700 μl, overlaid with 1200 μl of 30% (w/v), and 400 μl of 20% (w/v) iodixanol solution in assay buffer, centrifuged at 250.000 × g for 14 hours at 4°C in an SW60-Ti rotor (Beckman Coulter, Brea, CA). After centrifugation 10 equal fractions of 230 μl were collected from top, diluted 1:3 in assay buffer, and centrifuged at 100.000 × g for 2 h at 4°C in a TLA45/55 rotor (Beckman Coulter, Brea, CA). Vesicle pellets and semi-intact cells (input) were solubilised in SDS gel electrophoresis sample buffer and boiled at 95°C for 10 min prior to SDS-PAGE analysis.

### Electron microscopy

For electron microscopy (EM) of COPII vesicles reconstituted with GMP-PNP, gradient fraction 2 and 3 from three gradients were pooled, fixed with glutaraldehyde (2%) in assay, and subsequently pelleted by centrifugation at 100.000 × g for 1h in an TLA45/55 rotor (Beckman Coulter, Brea, CA). Resin-embedding and electron microscopy was carried out as described previously (Adolf et al., 2013).

### COPII vesicle formation for quantitative mass spectrometric analysis

For SILAC-based mass spectrometric analysis of COPII vesicles, HeLa cells ACC57 (DSMZ, Braunschweig, Germany) were cultivated at 37°C with 5% CO_2_ in DMEM medium containing and either natural amino acids or ^15^N_2_^13^C_6-_lysine (Lys-8) and ^15^N_4_^13^C_6_-arginine (Arg-10) (SILAC-Lys8-Arg10-Kit, Silantes, Munich, Germany) supplemented with 10% fetal bovine serum. Amino acid conversion of arginine to proline was prevented by addition of 200 μg/ml proline to the SILAC medium (Bendall et al., 2008). The COPII vesicle budding assay and density gradient isolation of vesicles were essentially carried out as described above, with the sole exception that after centrifugation the first 200 μl of the gradient were discarded, and a second fraction of 500 μl containing the vesicles was harvested. For a single label-switch experiment 4 × 6 budding reactions were carried out in parallel, mixed accordingly (e.g. vesicles from cells labelled with ‘light’ amino acids with control fractions from cells labelled with ‘heavy’ amino acids and vice versa). Mixed samples were run on a 10 well 1.5 mm 10% Tris-Glycine gel (Thermo Fisher Scientific, Waltham, MA) until the whole sample had entered the gel, fixed and stained with Roti®-Blue colloidal Coomassie (Carl Roth GmbH, Karlsruhe, Germany) and further processed for mass spectrometric analysis.

### Mass spectrometric analysis

Gel pieces were reduced with DTT, alkylated with iodoacetamid and digested with trypsin using the DigestPro MS platform (Intavis AG, Cologne, Germany) following the protocol described previously (Shevchenko et al., 2006). Peptides have then been analysed by liquid chromatography-mass spectrometry (LCMS) using an UltiMate 3000 LC (Thermo Scientific, Waltham, MA) coupled to either an Orbitrap Elite or a Q-Exactive mass spectrometer (Thermo Scientific, Waltham, MA). Peptides analysed by the Orbitrap Elite have been loaded on a C18 Acclaim PepMap100 trap-column (Thermo Fisher Scientific, Waltham, MA) with a flow rate of 30 μl/min 0.1% TFA. Peptides were eluted and separated on an C18 Acclaim PepMap RSLC analytical column (75 μm × 250 mm) with a flow rate of 300 nl/min in a 2 h gradient of 3% to 40% buffer B (0.1% formic acid, 90 % acetonitrile) in buffer A (0.1% formic acid, 1 % acetonitrile). MS data were acquired with an automatic switch between a full scan and up to 30 data-dependent MS/MS scans.

Peptides analysed on the Q-Exactive have been directly injected to an analytical column (75 μm × 300 mm), which was self-packed with 3 μm Reprosil Pur-AQ C18 material (Dr. Maisch HPLC GmbH, Ammerbruch-Entringen, Germany) and separated using the same gradient as described before. MS data were acquired with an automatic switch between a full scan and up to 15 data-dependent MS/MS scans.

Data analysis was carried out with MaxQuant version 1.5.3.8 (Cox and Mann, 2008) using standard settings for each instrument type and searched against a human specific database extracted from UniProt (UniProt Consortium). Carbamidomethylation of cysteine was specified as fixed modification; oxidation of methionine, deamidation of asparagine or glutamine and acetylation of protein N-termini was set as variable modification. ‘Requantify’ as well as ‘Match Between Runs’ options were both enabled. Results were filtered for a 1% false discovery rate (FDR) on peptide spectrum match (PSM) and protein level. MaxQuant output files have been further processed and filtered using self-compiled R-scripts and ribosomal, mitochondrial, and COPII coat proteins were manually removed to obtain the final datasets (SI Table 1 and 2).

### SNARE-GST binding assay

The SNARE-GST pull down assay was carried out essentially as described previously (Adolf et al., 2016). In brief, 150 μg of GST or 100 μg Syntaxin5^205-328^-GST fusion proteins were incubated with 20 μl of 50% (v/v) Glutathione Sepharose 4 fast flow (GE Healthcare, Little Chalfont, UK) bead-slurry in assay buffer (25 mM HEPES pH 7.2 (KOH), 150 mM KOAc, 2 mM MgOAc, 0.02% (v/v) monothioglycerol) in a total volume of 0.5 ml for 1h at 4°C. Glutathione Sepharose beads were washed three times with 1000 μl assay buffer and subsequently incubated with 30 μg of the indicated Sec23/24D complexes in assay buffer in a total volume of 500 μl for 1h at 4°C. Subsequently, Glutathione Sepharose beads were washed 3 times with 1000 μl assay buffer and bound Sec23/24D was eluted by boiling for 5 min at 95°C in SDS-PAGE sample buffer. Input fractions of the Sec23A/24D complexes (2%) and 40% of samples were analysed by SDS–PAGE and Coomassie brilliant blue (CBB) staining and Western blotting.

## Acknowledgements

We are grateful to all members of the Wieland lab for critical discussions, comments, and support. We are grateful to Hilmar Bading, Department of Neurobiology and Interdisciplinary Center for Neurosciences, University of Heidelberg, for providing the opportunity to carry out the electron microscopy work in his laboratory. This work was supported by the German Research Council, Grant SFB 638, A10, and DFG-Einzelprojekt Wi-654/12-1 to FTW.

## Author contributions

FA and FTW conceived and designed experiments; FA cloned, expressed and purified the recombinant proteins and performed most reconstitution experiments for MS and EM analysis; MR assisted with sample preparation for MS data acquisition and performed reconstitution experiments with Sec24D mutants; BH preformed MS analysis; AH preformed EM; FA and FTW analysed the data and wrote the manuscript.

## SI Figure Legends

**SI Fig.1:**
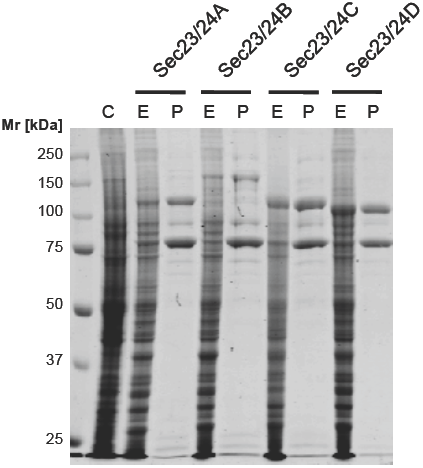
Expression and purification of recombinant mammalian Sec23/24 coat subcomplexes in Sf9 insect cells. The four mammalian Sec24 isoforms were expressed in complexes with Sec23A in Sf9 insect cells (E). Cells were harvested 72h post infection and protein complexes were purified by affinity purification and gel filtration (P). Equal volumes of non-infected control cells (C), and Sf9 cells expressiing His6-Sec23A/24A, His6-Sec23A/23B, His6-Sec23A/23C, and His6-Sec23A/23D, as well as 2 μg of the respective purified COPII inner coat subcomplex were separated by SDS polyacrylamide gel electrophoresis and stained with Coomassie brilliant blue (CBB).

**SI Fig.2:**
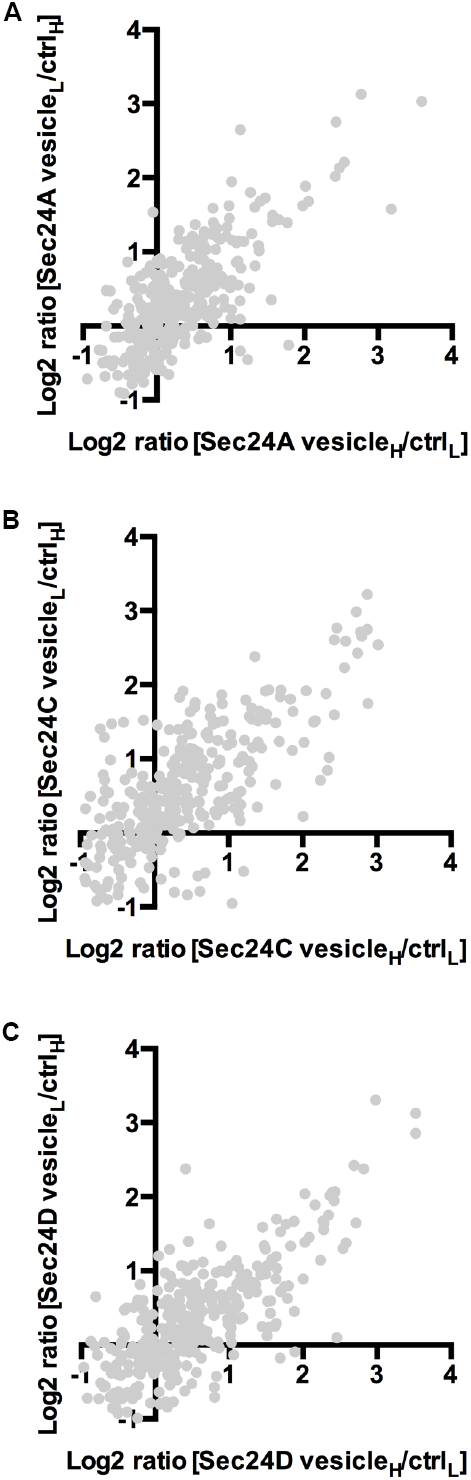
Proteomics of *in vitro* reconstituted isotopic COPII vesicles. Scatter plot depicting all quantified proteins in SILAC label-switch experiments of COPII vesicles reconstitutions from SIC labelled with ‘light’ or ‘heavy’ amino acids (A-C) with the isoforms Sec23A/24A (A), Sec23A/24C (B), or Sec23A/24D (C).

